# Catalytic inhibition of H3K9me2 writers disturbs epigenetic marks during bovine nuclear reprogramming

**DOI:** 10.1101/847210

**Authors:** RV Sampaio, JR Sangalli, THC De Bem, DR Ambrizi, M del Collado, A Bridi, ACFCM Ávila, CH Macabelli, LJ Oliveira, JC da Silveira, MR Chiaratti, F Perecin, FF Bressan, LC Smith, PJ Ross, FV Meirelles

**Author notes:** **Correspondence** Rafael Vilar Sampaio, Department of Veterinary Medicine, Faculty of Food Engineering and Animal Science, University of Sao Paulo, Pirassununga, Sao Paulo, Brazil.

## Abstract

Orchestrated events, including extensive changes in epigenetic marks, allow a somatic nucleus to become totipotent after transfer into an oocyte, a process termed nuclear reprogramming. Recently, several strategies have been applied in order to improve reprogramming efficiency, mainly focused on removing repressive epigenetic marks such as histone methylation from the somatic nucleus. Herein we used the specific and non-toxic chemical probe UNC0638 to inhibit the catalytic activity of the histone metyltransferases EHMT1 and EHMT2. Either the donor cell (before reconstruction) or the early embryo was exposed to the probe to assess its effect on developmental rates and epigenetic marks. First, we showed that the treatment of bovine fibroblasts with UNC0638 did mitigate the levels of H3K9me2. Moreover, H3K9me2 levels were decreased in cloned embryos regardless of treating either donor cells or early embryos with UNC0638. Additional epigenetic marks such as H3K9me3, 5mC, and 5hmC were also affected by the UNC0638 treatment. Therefore, the use of UNC0638 did diminish the levels of H3K9me2 and H3K9me3 in SCNT-derived blastocysts, but this was unable to improve their preimplantation development. These results indicate that the specific reduction of H3K9me2 by inhibiting EHMT1/2 causes diverse modifications to the chromatin during early development, suggesting an intense epigenetic crosstalk during nuclear reprogramming.

## INTRODUCTION

Cloning by somatic cell nuclear transfer (SCNT) is an inefficient technique largely because of incomplete nuclear reprogramming. The persistent epigenetic memory from the somatic cell nucleus is considered one of the main barriers for an efficient reprogramming process ^1^. In most cases, the recipient oocyte used in SCNT fails to erase residual epigenetic marks from somatic nucleus, which are retained in the embryo and lead to abnormal gene expression ^2^. For instance, when compared to in vitro fertilized (IVF) embryos, SCNT embryos show an aberrant pattern of gene expression during the period of embryonic genome activation (EGA) ^3,4^. According to previous works, aberrant gene expression in cloned embryos is linked with altered DNA and histone demethylation ^5,6^. Hence, epigenetic errors might lead to problems such as abortion, placental and fetal abnormalities, among others ^7^.

One of the main anomalies identified during SCNT development relates to its aberrant EGA ^3,4,8^. Recently, genomic areas refractory to reprogramming were identified and named as “reprogramming resistant regions” (RRR) ^8^. In cattle, major EGA occurs at the 8/16-cell stage ^9^ and it is possible that augmented levels of H3K9me2 and H3K9me3 during this stage are responsible for the poor development of SCNT-derived embryos in cattle ^6,10^.

Modification on the lysine residue at the position 9 on histone H3 is one of the most studied epigenetic marks. EHMT2 and the related molecule EHMT1 (also known as G9a and GLP) are the main enzymes responsible for methylation at this site ^11^. EHMT1/2 are SET (Su(var)3-9, E(z), and Trithorax) domain-containing proteins, a conserved domain present in other epigenetic enzymes that uses the cofactor S-adenosyl-l-methionine (SAM) to achieve methylation on histones and other proteins ^12^. They also have an Ankyrin (ANK) repeat domain, which can generate and read the same epigenetic mark and are considered as a motif to interact with other proteins ^13^. There is a well-known crosstalk between DNA methylation and histone modifications. DNA methyltransferase DNMT1 directly binds EHMT2 to coordinate their modification during DNA replication ^14^. EHMT2 is also implicated in genomic imprinting regulation ^15,16^ and its absence results in the impairment of placenta-specific imprinting ^17^. However, the enzymatic activity of EHMT1/2 has recently been shown to be dispensable for imprinted DNA methylation, since EHMT1/2 can recruit DNMTs via its ANK domain ^18^.

Small molecules have been used to inhibit EHMT1/2 in an attempt to reduce H3K9me2 and facilitate nuclear reprogramming during induced pluripotent stem cell (iPSC) generation ^19,20^ and SCNT ^21,22^; however, even with the intense crosstalk between histone modification and DNA methylation, the effects of the inhibition of H3K9me2 writers on DNA methylation and other epigenetic marks remain unknown. In this study, we hypothesized that the inhibition of EHMT1/2 activity either before or after nuclear results in decreased H3K9me2 levels and improves development of SCNT-derived embryos. Additionally, we investigated throughout preimplantation development the consequences of EHMT1/2 inhibition on additional epigenetic marks and expression of genes related to epigenetic regulation.

## RESULTS

### EHMT1/2 inhibition reduces H3K9me2 levels in donor cells

To test whether the UNC0638 treatment would be suitable for the bovine species, fetal fibroblasts were treated with 250 nM of UNC0638 for 96 h and the levels of H3K9me2 were analyzed by immunofluorescence (IF) and western blot (WB). Both IF and WB results showed that UNC0638 treatment decreased H3K9me2 levels in bovine fibroblasts when compared to non-treated controls (Fig.1A-D). Moreover, the treatment with UNC0638 did not impact on the levels of H3K9me3, validating the efficiency and specificity of UNC0638 as a bovine EHMT1/2 inhibitor.

Next, we aimed at investigating whether the lower levels of H3K9me2 were kept after nuclear transfer. Hence, these fibroblasts were used as nuclear donors. In agreement with the result found in treated fibroblasts (Figure 1), the use of these cells as nuclear donors revealed that lower levels of H3K9me2 were present in 1-cell embryos 18 h after oocyte reconstruction. This gives evidence that the chromatin remodeling occurring immediately after nuclear transfer was not able to revert the treatment effect on H3K9me2 levels (Figure 2). To further evaluate the implications of the treatment, we also assessed the levels of H3K9me3, 5mC and 5hmC in 1-cell embryos. Although the levels of 5mC and 5hmC remained unchanged, increased levels of H3K9me3 were found after 18 h of oocyte reconstruction (Figure 2). Given that the levels of H3K9me3 were unaltered in fibroblasts prior to nuclear transfer (Figure 1), it is likely that this change occurred after oocyte reconstruction.

**Figure 1.**
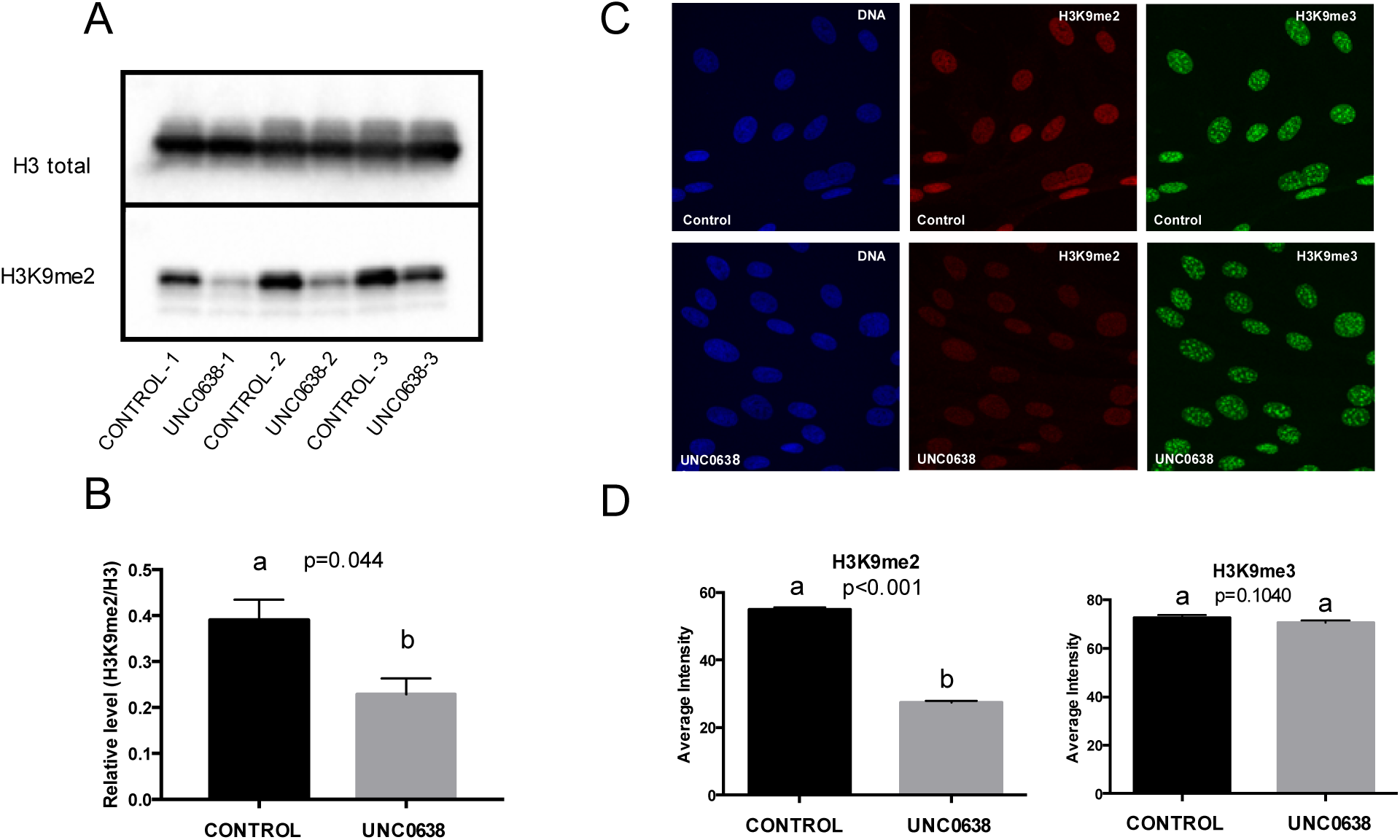
UNC0638 reduces the abundance of the H3K9me2 in bovine fibroblasts. (**A**) Western blots and (**B**) quantification of H3K9me2 of fetal fibroblasts treated with either 0.01% of DMSO (CONTROL) or 250 nM of UNC0638 (UNC) for 96 h. Results from three pairs of biological replicates are shown. H3K9me2 levels were calculated in relation to total histone 3 (H3). Western blot images were cropped for illustrative purposes and images of full blots are presented in Supplemental Fig.1. (**C**) Levels of H3K9me2 and H3K9me3 determined by immunofluorescence (IF). Representative images of bovine fetal fibroblasts treated for 96 h with 0.01% DMSO (Control) or 250nM UNC0638 (UNC0638). Images show fibroblasts nuclei stained with DAPI (left column), anti-H3K9me2 (red; middle column), and anti-H3K9me3 (green; right column) antibodies. Images were taken using the same magnification (63x) and laser power, thereby enabling direct comparison of signal intensities. (**D**) Quantification of H3K9me2 and H3K9me3 fluorescence intensity in nuclei of fibroblasts treated or not with UNC0638 (UNC). Values are presented as mean ± S.E.M. and different letters indicate significant differences (p<0.05).

**Figure 2.**
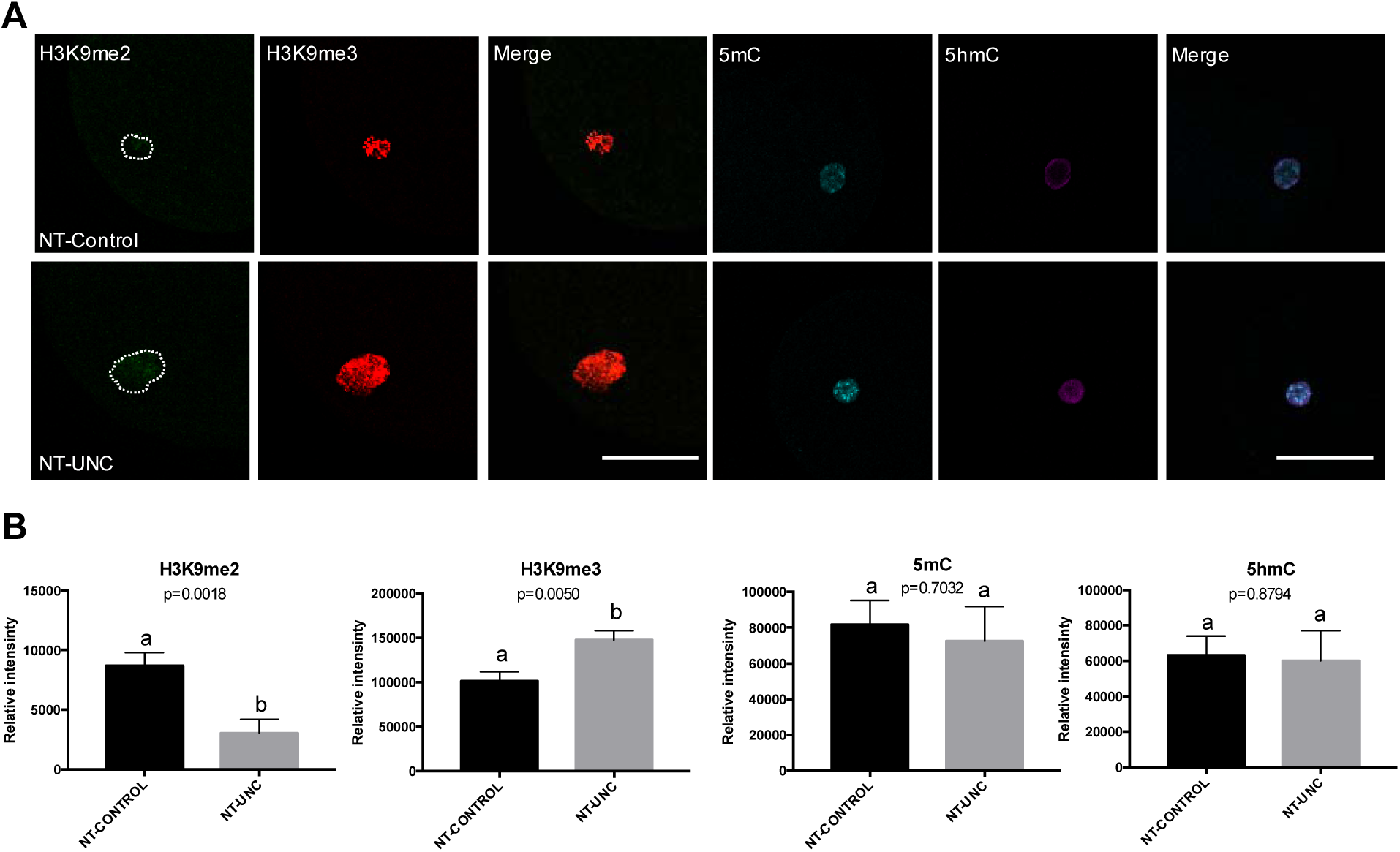
Levels of H3K9me2, H3K9me3, 5mC, and 5hmC in SCNT 1-cell embryos derived from nuclear donor cells treated with UNC0638. (**A**) One-cell embryos derived from control donor cells are displayed on first lane and zygotes derived from donor cells treated with UNC0638 are displayed on second lane. Columns 1, 2, 4, and 5 show 1-cell embryos stained with antibody anti-H3K9me2 (green), anti-H3K9me3 (red), anti-5mC (cyan), and anti-5hmC (magenta), respectively. Merge images from H3K9me2+H3K9me3 and 5mC+5hmC are displayed in columns 3 and 6, respectively. All images were taken at the same magnification and at the same laser power, thereby enabling direct comparison of signal intensities. Scale bar 100 μm. (**B**) Quantification of H3K9me2, H3K9me3, 5mC, and 5hmC levels in SCNT embryos derived from donor cells treated or not with UNC0638. Embryos nuclei from 3 different biological replicates were analyzed to investigate the levels of H3K9me2 + H3K9me3 (NT-Control N=13; NT-UNC N=10) and 5mC + 5hmC (NT-Control N=9; NT-UNC0638 N=10). Data are presented as mean ± S.E.M. Different letters above bars indicate significantly differences (p<0.05). NT-Control: Zygotes derived from cells treated with 0.01% of DMSO. NT-UNC: Zygotes derived from cells treated with 250nM of UNC0638.

Similar to what was seen in 1-cell embryos, the levels of H3K9m2 were decreased in both 8/16-cell embryos and blastocysts (Figure 3 and 4) produced using donor cells treated with UNC0638. This result supports the idea that the low levels of H3K9m2 presented in fibroblasts were kept after nuclear transfer and up to the blastocyst stage. In contrast to the 1-cell stage, the levels of H3K9me3 were no longer altered in 8/16-cell embryos derived from UNC0638 treated cells (Figure 3). Furthermore, lower levels of H3K9me3 were found in these embryo at the blastocyst stage (Figure 4). With respect to 5mC and 5hmC, the levels of these marks were increased in 8/16-cell stage embryos reconstructed using UNC0638-treated cells (Figure 3), but not altered in blastocysts (Figure 4). This indicates a complex and dynamic epigenetic effect of the UNC0638 treatment on DNA methylation during early development.

**Figure 3.**
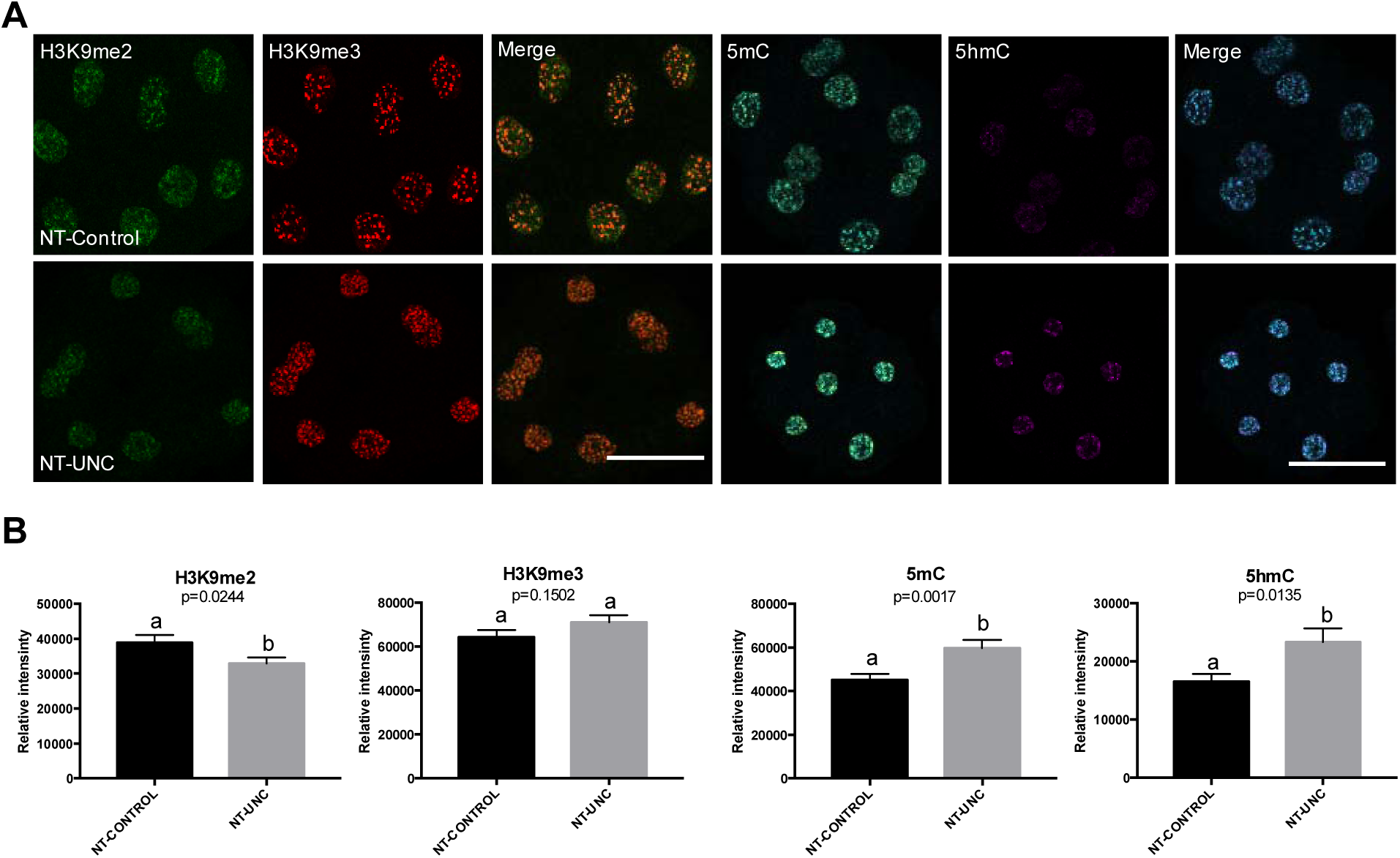
Levels of H3K9me2, H3K9me3, 5mC, and 5hmC in 8/16-cells embryos derived from nuclear donors previous treated with UNC0638. (**A**) Embryos at 8/16-cells stage derived from nuclear donors control are displayed on first lane and embryos derived from cells treated with UNC0638 are displayed on second lane. Columns 1, 2, 4, and 5 are showing embryos stained with antibody anti-H3K9me2 (green), anti-H3K9me3 (red), anti-5mC (cyan), and anti-5hmC (magenta) are displayed in columns 3 and 6, respectively. All images were taken at the same magnification and at the same laser power, thereby enabling direct comparison of signal intensities. Scale bar 100 μm. (**B**) Quantification of H3K9me2, H3K9me3, 5mC, and 5hmC levels in SCNT embryos derived from cells treated or not with UNC0638. Embryos nuclei from 3 different biological replicates were analyzed to investigate the levels of H3K9me2 + H3K9me3 (NT-Control N=10; NT-UNC N=8) and 5mC + 5hmC (NT-Control N=8; NT-UNC0638 N=9). Data as presented as mean ± S.E.M, and bars with different letters are significantly different p<0.05). NT-Control: 8/16-cells embryos derived from cells treated with 0.01% of DMSO. NT-UNC: 8/16-cells embryos derived from cells treated with 250nM of UNC0638.

**Figure 4.**
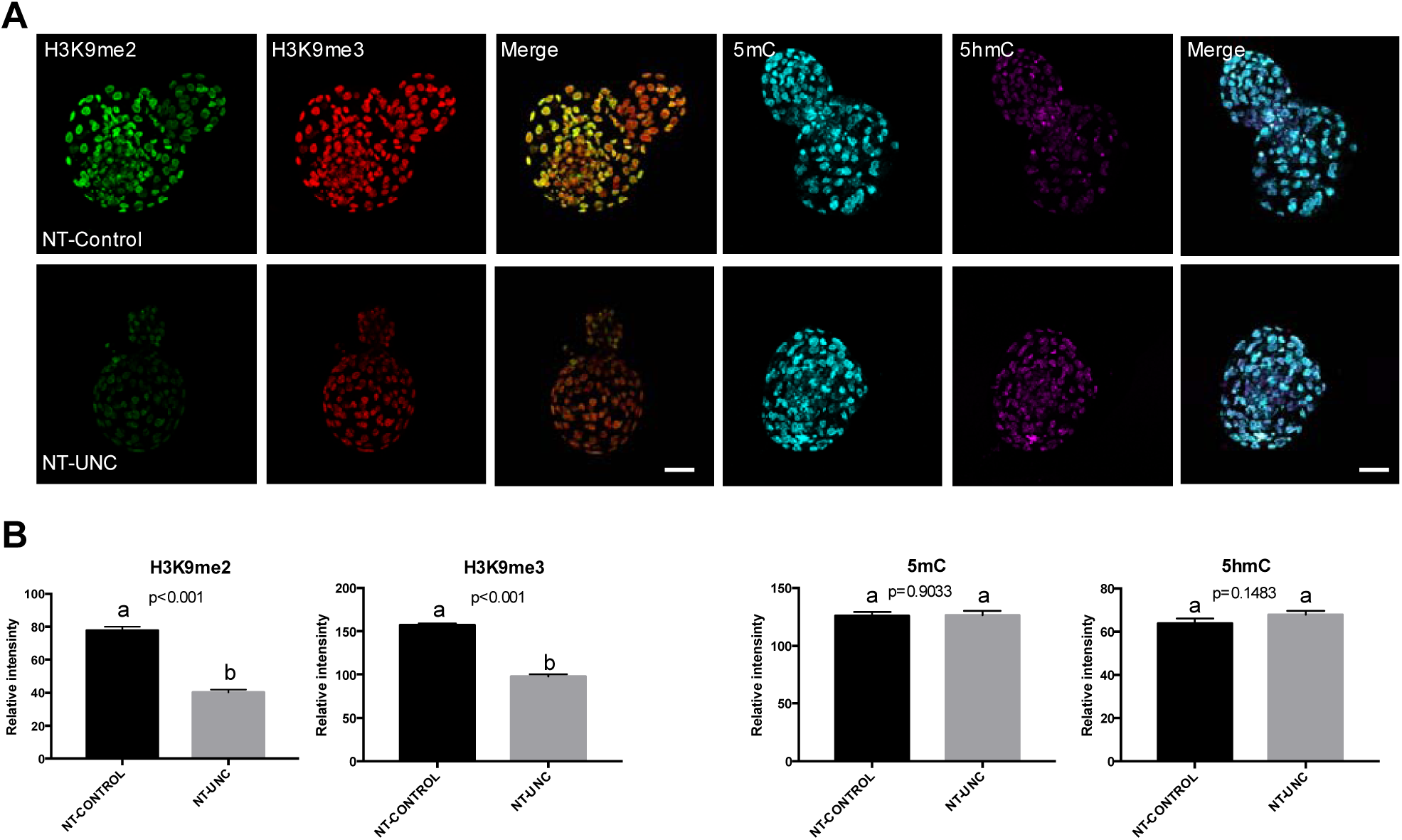
Levels of H3K9me2, H3K9me3, 5mC, and 5hmC in blastocysts derived from nuclear donor cells treated with UNC0638. (**A**) Blastocysts derived from nuclear donor controls are displayed in first lane and embryos derived from cells treated with UNC0638 are displayed in second lane. Columns 1, 2, 4, and 5 are showing embryos stained with antibody anti-H3K9me2 (green), anti-H3K9me3 (red), anti-5mC (cyan), and anti-5hmC (magenta), respectively. Merge images from H3K9me2+H3K9me3 and 5mC+5hmC are displayed in columns 3 and 6, respectively. All images were taken at the same magnification and at the same laser power, thereby enabling direct comparison of signal intensities. Scale bar 100 μm. (**B**) Quantification of H3K9me2, H3K9me3, 5mC, and 5hmC levels in SCNT embryos derived from cells treated or not with UNC0638. Embryos nuclei from 3 different biological replicates were analyzed to investigate the levels of H3K9me2 + H3K9me3 (NT-Control N=15; NT-UNC N=12) and 5mC + 5hmC (NT-Control N=11; NT-UNC0638 N=13). Data as presented as the mean ± S.E.M. and means with different letters are significantly different (p<0.05). NT-Control: Blastocysts derived from cells treated with 0.01% of DMSO. NT-UNC: Blastocyst derived from cells treated with 250nM of UNC0638.

In cattle, the high levels of H3K9me2 present in somatic nuclei persist during development of SCNT-derived embryos and might play a role as a barrier for nuclear reprogramming ^6,23^. Given the success of the UNC0638 treatment in decreasing the levels of H3K9me2 in fibroblasts and SCNT-derived embryos, we assessed its impact on developmental rates. Remarkably, no effect was observed on both cleavage (*p* = 0.33) and blastocyst (*p* = 0.73) rates (Table 1). Therefore, our findings suggest that the levels of H3K9me2 present in donor cells are not linked with the preimplantation development of SCNT embryos.

**Table 1.** Development of SCNT zygotes using nuclear donor cells treated with the EHMT1/2 inhibitor UNC0638 before oocyte reconstruction.

| Treatment | Cultured zygotes | Cleavage ± S.E.M % | Blastocyst % ± S.E.M |
| --- | --- | --- | --- |
| NT-CONTROL | 287 | 70.89 ± 5.6 | 26.95 ± 5.5 |
| NT-UNC | 300 | 62.18 ± 6.6 | 24.31 ± 5.2 |

Note: Development rates were obtained from the number of embryos that cleaved or reached the blastocyst stage per reconstructed oocytes (cultured zygotes). Results from 9 replicates.

### EHMT1/2 inhibition in SCNT embryos affects epigenetic marks during early development

In agreement with the above findings, culture of cloned embryos for 55 h in the presence of UNC0638 (from oocyte reconstruction to the 8/16-cell stage) also led to decreased levels of H3K9me2 and increased levels of H3K9me3, 5mC and 5hmC in 8/16-cell stage embryos (Figure 5). In this experiment donor cells were not treated, providing evidence that the treatment after oocyte reconstruction was also effective. Moreover, the levels of both H3K9me2 and H3K9me3 were decreased at the blastocyst stage in comparison with untreated embryos (Figure 6). This reversal of H3K9me3 from higher to lower levels is consistent with that observed when the treatment was performed in donor cells (Figure 2 to 4). Similarly, decreased levels of 5hmC were present in blastocysts derived from the UNC0638 treatment (Figure 6), suggesting that UNC0638 results in dynamic changes on the levels of H3K9me3, 5mC and 5hmC (Figure 3). Overall, in spite of slight differences, treatment of either donor cells or early embryos with UNC0638 resulted in a similar pattern of epigenetic modification.

**Figure 5.**
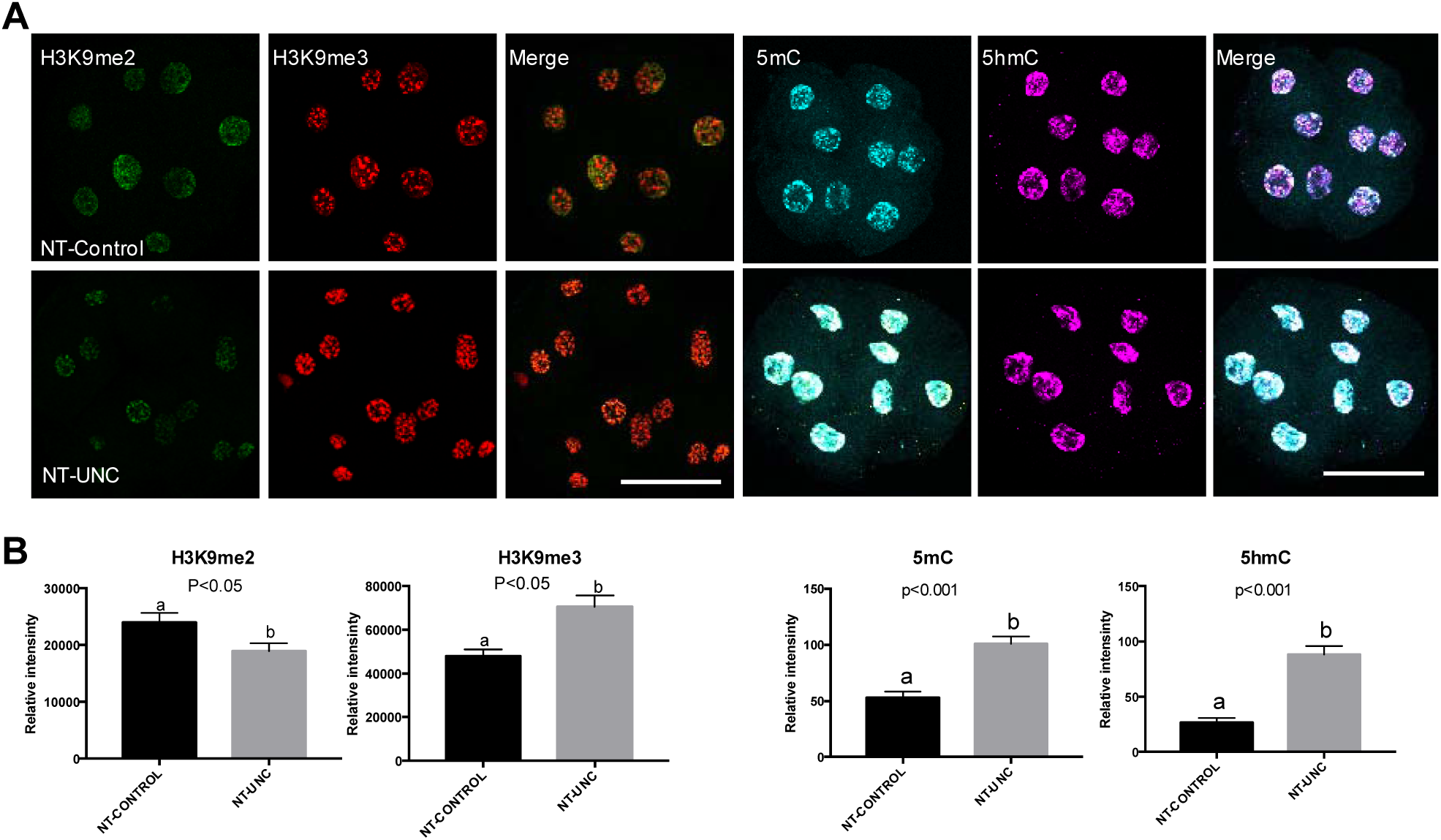
Levels of H3K9me2, H3K9me3, 5mC, and 5hmC in 8/16-cells embryos treated with UNC0638. (**A**) Embryos at 8/16-cell stage controls are displayed in first lane and embryos previous treated with UNC0638 are displayed in second lane. Columns 1, 2, 4, and 5 are showing embryos stained with antibody anti-H3K9me2 (green), anti-H3K9me3 (red), anti-5mC (cyan), and anti-5hmC (magenta), respectively. Merge images from H3K9me2+H3K9me3 and 5mC+5hmC are displayed in columns 3 and 6, respectively. All images were taken at the same magnification and at the same laser power, thereby enabling direct comparison of signal intensities. Scale bar 100 μm. (**B**) Quantification of H3K9me2, H3K9me3, 5mC, and 5hmC levels in SCNT embryos treated or not with UNC0638. Embryos nuclei from 3 different biological replicates were analyzed to investigate the levels of H3K9me2 + H3K9me3 (NT-Control N=8; NT-UNC N=8) and 5mC + 5hmC (NT-Control N=8; NT-UNC0638 N=7). Data are presented as mean ± S.E.M. Bars with different letters are significantly different (p<0.05). NT-Control: 8/16-cells embryos previous treated with 0.01% of DMSO. NT-UNC: 8/16-cell embryos previous treated with 250nM of UNC0638.

**Figure 6.**
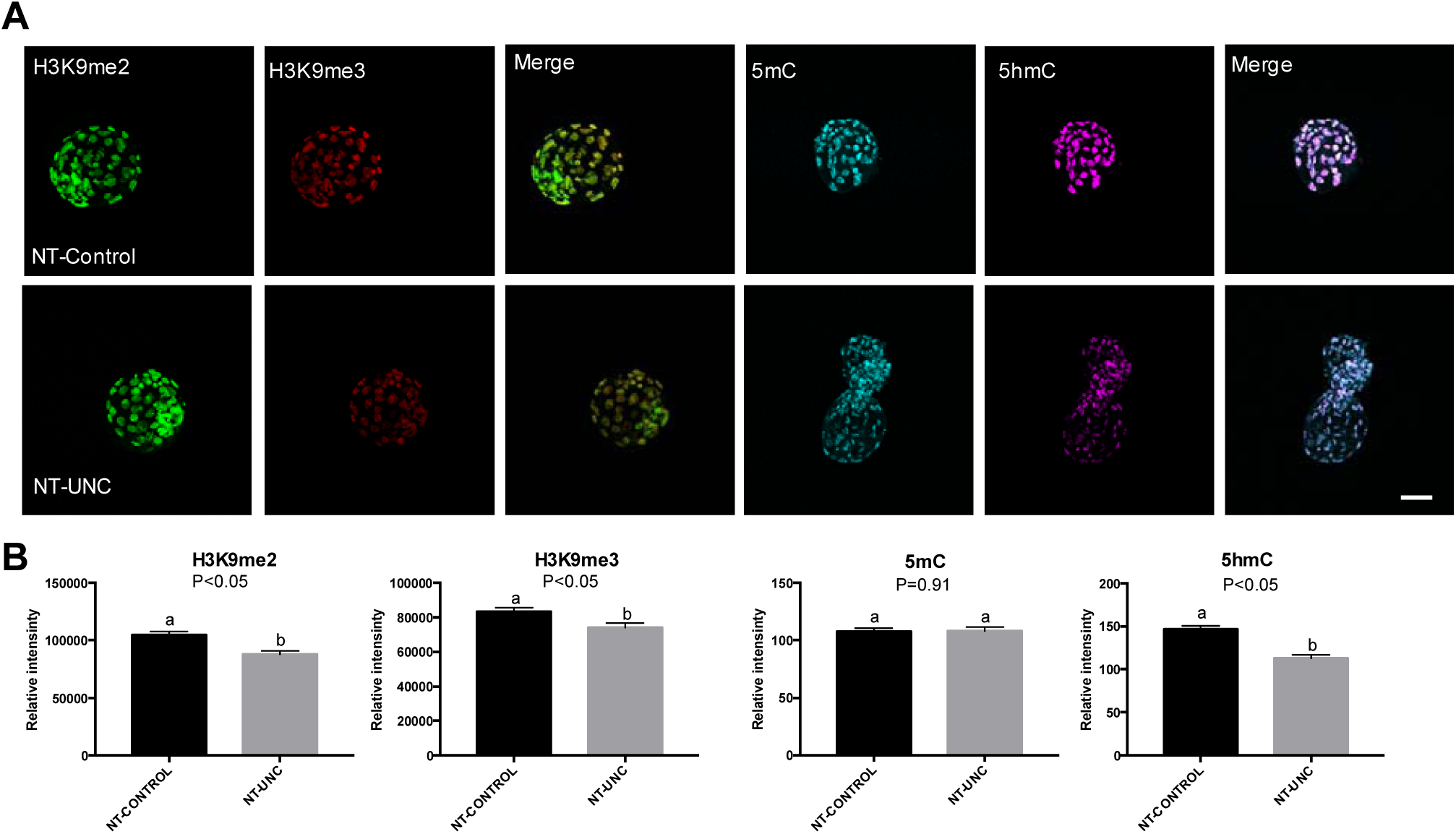
Levels of H3K9me2, H3K9me3, 5mC, and 5hmC in blastocysts treated with UNC0638. **(A)** Embryos at 8/16-cell stage controls are displayed in first lane and embryos previous treated with UNC0638 are displayed in second lane. Columns 1, 2, 4, and 5 are showing embryos stained with antibody anti-H3K9me2 (green), anti-H3K9me3 (red), anti-5mC (cyan), and anti-5hmC (magenta), respectively. Merge images from H3K9me2+H3K9me3 and 5mC+5hmC are displayed in columns 3 and 6, respectively. All images were taken at the same magnification and at the same laser power, thereby enabling direct comparison of signal intensities. Scale bar 100 μm. **(B)** Quantification of H3K9me2, H3K9me3, 5mC, and 5hmC levels in SCNT embryos treated or not with UNC0638. Embryos nuclei from 3 different biological replicates were analyzed to investigate the levels of H3K9me2 + H3K9me3 (NT-Control N=10; NT-UNC N=7) and 5mC + 5hmC (NT-Control N=7; NT-UNC0638 N=8). Data are presented the mean ± S.E.M. Bars with different letters are significantly different (p<0.05). NT-Control: Blastocysts derived from embryos previous treated with 0.01% of DMSO. NT-UNC: Blastocysts derived from embryos previous treated with 250nM of UNC0638.

Next, we sought to determine the effect of the UNC0638 treatment on transcript levels of i) the main inhibited targets, such as *EHMT1* and *EHMT2*; ii) genes potentially related to the activity of EHMTs, such as *DNMT1, DNMT3A*, and *DNMT3B* ^14,24^; and, iii) genes linked with the role played by H3K9me2 during embryonic development, such as *TET1, TET2*, and *TET3* ^25^. However, no effect on transcript abundance was found when considering 8/16-cell embryos and blastocysts (Figure 7). These results indicate that the changes observed on epigenetic marks were independent of gene expression, i.e. at a post-transcriptional level.

**Figure 7.**
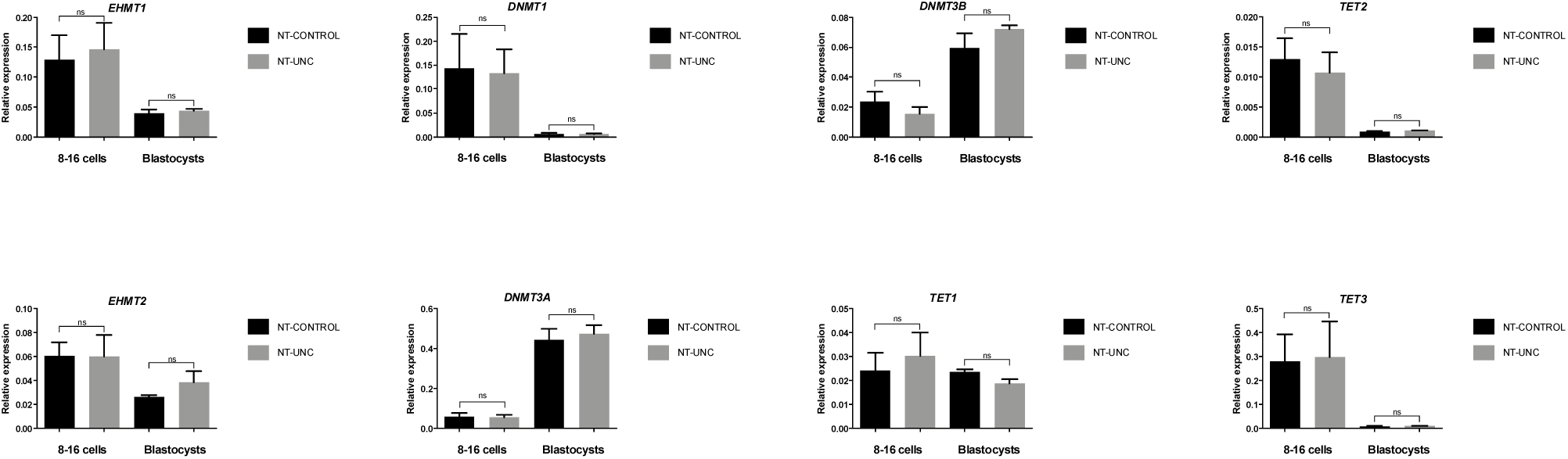
Treatment with UNC0638 SCNT embryos during early development does not change transcript levels of epigenetic writer and eraser genes at 8/16-cell and blastocyst stages. The mRNA levels of *EHMT1, EHMT2, DNMT1, DNMT3A, DNMT3B, TET1, TET2*, and *TET3* were quantified using real-time RT-PCR of SCNT embryos treated or not with 250nM of UNC0638. Pool of 5 embryos from 3 different biological replicates (total 15 per group) were analyzed to investigate the levels. Transcript quantities were normalized to the geometric mean of two housekeeping genes (PPIA and RPL15). Data are shown as relative expression, and the values as the mean ± S.E.M. (p > 0.05).

Finally, treatment of SCNT-derived embryos with UNC0638 did not impact on both cleavage (*p* = 0.65) and blastocyst (*p* = 0.13) rates (Table 2).

**Table 2.** Development of SCNT-reconstructed oocytes treated with the EHMT1/2 inhibitor UNC0638 from activation to the 8/16-cell stage.

| Treatment | Cultured zygotes | Cleavage ± S.E.M % | Blastocyst % ± S.E.M |
| --- | --- | --- | --- |
| NT-CONTROL | 249 | 83.76 ± 2.1 | 29.34 ± 3.4 |
| NT-UNC | 255 | 82.06 ± 3.0 | 35.83 ± 2.3 |

Note: Development rates were obtained from the number of embryos that cleaved or reached the blastocyst stage per reconstructed oocytes placed in culture (cultured zygotes). Results from 10 replicates.

## DISCUSSION

In summary, our findings indicate that the treatment with UNC0638 was able to reduce the levels of H3K9me2 regardless of treating either donor cells or the early embryo derived from SCNT. Importantly, the lower levels of H3K9me2 were maintained throughout the preimplantation development, but this failed to enhance developmental rates. Additionally, the levels of H3K9me3, 5mC and 5hmC in SCNT-derived embryos were impacted by the UNC0638 treatment, but not the transcriptional activity of genes related to the role of EHMTs and H3K9me2.

Residual epigenetic memory has been widely considered responsible for improper reprogramming of somatic cells ^26^. The high levels of H3K9me2 present in somatic nuclei persist during the development of bovine SCNT embryos ^6,23^. Aiming to decrease H3K9me2 levels, we used UNC0638, a small molecule that specifically inhibits the catalytic activity of histone methyltransferases EHMT1/2 in a non-toxic manner as previously shown in mice and sheeps ^27,28^. Likewise, our treatment proved effective, resulting in lower levels of H3K9me2 in somatic cells exposed to UNC0638. This reduction was observed throughout embryogenesis of SCNT embryos, regardless of the moment of the treatment (i.e. in donor cells or in early embryos). We then hypothesized that distinct time of expression between H3K9me2 writers and erasers might explain the lower levels of H3K9me2 since the expression of *EHMT1* and *EHMT2* in bovine is higher in oocytes than in early embryos ^29^ while the expression of lysine demethylases is increased later in development ^30^. Herein, we found no effect of the UNC0678 treatment when main regulatory genes were analyzed (Figure 7). This result is consistent with a previous report with somatic cells in which the UNC0638 treatment did not affect neither the protein levels of EHMT1 or EHMT2 nor the mRNA levels of *EHMT2*, indicating that changes in H3K9me2 were a consequence of decreased enzymatic activity ^28^.

As expected, different epigenetic marks were also found to be changed herein by EHMT1/2 inhibition. During the earliest stages of embryogenesis, we observed an increase in H3K9me3 levels, which was followed by a drop in blastocysts. This increase in H3K9me3 has been reported as a classical epigenetic alteration secondary to the low levels of H3K9me2 due the preferential activity of SUV39H for mono or di-methylated H3K9 as a substrate to form H3K9me3 ^31,32^. In fact, demand for substrate along with demethylase activity is a reasonable explanation for the subsequent decline in H3K9me3 levels as H3K9me2 is required for H3K9me3 formation. In agreement with this, treatment with an EHMT2 inhibitor led to a decrease in H3K9me3 at target genes ^33^. Moreover, common genes were upregulated by the knockdown of *EHMT2* or *SUV39H1* ^34^, suggesting they share similar targets. However, given that H3K9me3 was not changed in somatic cells, these dynamic changes in H3K9me3 levels likely require methylation writers and erasers present in the early embryo ^35^.

Regarding DNA modifications, analysis of SCNT embryos revealed an increase at 8/16-cell stage in 5mC and 5hmC levels, regardless of the moment the treatment was performed (Figure 2-4). Due to the fact that 5mC is required for 5hmC formation ^36,37^, we speculate that the increase in 5hmC was likely a consequence of increased 5mC availability. Actually, we expected an opposite result given that knockout of EHMT1/2 in ESCs leads to global DNA demethylation ^38,39^. This is in agreement with the idea that the presence of H3K9me2 in the maternal pronucleus, together with DPPA3/STELLA, prevents active DNA demethylation by Tet-family enzymes ^40^. Nonetheless, it was recently shown, both in IVF and SCNT embryos, that DNA demethylation may be independent of H3K9me2; only DPPA3/STELLA was needed ^41^. Moreover, a previous work showed that the treatment with an EHMT2 inhibitor not only decreased H3K9me2, but also increased DNA methylation in neuroblastoma cells ^42^. It might be possible that herein inhibition of EHMT1/2 catalytic activity prompted EHMT1/2 to cooperate in additional functions. In fact, the involvement of EHMT2 in DNA methylation goes beyond its methylase activity ^24,38^. However, the process involved in this offset mechanism remains elusive. The DNA hypermethylation of cloned embryos is one of the major failures of proper chromatin remodeling during nuclear reprogramming ^43,44^, however, in contrast to changes at a global level, severe demethylation in important regions as satellite I and imprinted genes ^43,44^ could also be responsible for such failure.

Finally, its noteworthy that the unaltered transcript levels of UNC0638-treated embryos suggest a very specific effect of the treatment on EHMT1/2 inhibition. This is consistent with the finding that treatment with UNC0638 does not affect developmental rates as reported here and previously in mice ^45^, goats ^46^, and sheep ^27,47^. It is likely the effect of the UNC0638 is species-specific as only in pigs it was shown to improve development of SCNT-derived embryos ^21,48^. Likewise, one might expect the UNC0638 treatment to improve developmental rates as it significantly mitigated the levels of H3K9me3 in blastocysts; this mark is considered a main epigenetic barrier for nuclear reprogramming. First described in mice, the RRRs are H3K9me3-enriched regions refractory to transcriptional activation in 2-cell cloned embryos, the moment when EGA occurs in mice ^8^. Similarly, reduction of H3K9me3 levels during bovine EGA improved SCNT efficiency ^10^, indicating that in this species, 8/16-cells stage may be the best period to test H3K9me3 reduction-based approaches. Taken together, these data suggest that EHMT1/2 contribute to the stability of different epigenetic marks during bovine reprogramming independently of their catalytic activity.

## CONCLUSION

Using a specific probe to inhibit the catalytic activity of EHMT1/2, we were able to mitigate the levels of H3K9me2 throughout early embryonic development. Inhibition, either in nuclear donor cells or during early development of SCNT-derived embryos triggered changes in H3K9me3, 5mC, and 5hmC levels independently of transcriptional changes. This suggests an intense crosstalk among histone modifications and DNA methylation during nuclear reprogramming in cattle. Nonetheless, although lower levels of H3K9me2 and H3K9me3 were achieved at the blastocyst stage, EHMT1/2 inhibition did not increase developmental rates. Therefore, the aberrant epigenetic remodeling present in SCNT-derived embryos may not be amended by inhibiting the catalytic activity of these epigenetic writers.

## METHODS

All chemicals and reagents used were purchased from Sigma-Aldrich Chemical Company (St. Louis, MO, USA), unless otherwise stated. The present study was approved by the ethical committee for the use of animals of the School of Veterinary Medicine and Animal Science of the University of Sao Paulo, protocol number 9076280114, and in agreement with the ethical principles of animal research. We adopted the International Guiding Principles for Biomedical Research Involving Animals (Society for the Study of Reproduction) as well.

### Cell culture

Fibroblast cell line was obtained from a 55-days-old crossbred (Gir x Holstein; *Bos indicus* x *Bos taurus*) fetus, as described previously ^49^. Concisely, a skin biopsy (1 cm^2^) was minced into small pieces with a scalpel blade, followed by digestion with 0.1% (w/v) collagenase for 3 h at 38.5 °C. The digested product was centrifuged at 300 × *g* for 5 min, and the pellet was resuspended in culture medium (α-MEM [GIBCO BRL, Grand Island, NY, USA] supplemented with 10% [v/v] fetal bovine serum [FBS] and 50 μg/ml gentamicin sulfate). Then, plated in 35-mm Petri dishes and cultured for 6 days in an incubator to establish the primary culture. After achievement of 70% of confluence (approximately 4×10^5^ cells), cells were detached from the plate using Tryple Express (GIBCO BRL); resuspended in culture medium supplemented with 10% of dimethyl sulfoxide (DMSO), and 50 μg/mL gentamicin sulfate; placed in cryotubes; and stored in liquid N_2_ until use.

### Cell treatment with UNC0638

For nuclear donor treatment, primary fetal fibroblasts were thawed and cultured as described above. After 24 h, culture medium was replaced with fresh medium containing 250nM of UNC0638, best concentration described previously ^28^. Forty-eight hours later, UNC0638 treatment was renewed for an additional 48h under serum starvation (culture media supplemented with 0.5% of FBS instead of 10%) to arrest the cell cycle at the G1/G0 stages. Dimethyl sulfoxide (DMSO) at same concentration of UNC0638 dilution (0.01%) and submitted to same culture conditions was used as control. Nighty-six hours after starting treatment (48h + 48h), cells were then used for immunostaining, western blotting analysis or SCNT. While for embryo treatment, nuclear donors were cultured for 48h under starvation medium without treatment.

### Immunostaining of somatic cells

For H3K9me2 and H3K9me3 analysis, we procedure as described previously ^50,51^ with few modifications. Cells were plated on coverslips and treated as described above. Then, cells were fixed with 4% paraformaldehyde for 15 min and permeabilized with D-PBS + 1% Triton X-100 for 10min and blocked for 30 min in D-PBS + 0.3%Triton X-100 + 1% BSA, were placed in primary antibody solution consisting of blocking buffer, a mouse antibody anti-H3K9me2 - Abcam (ab1220, 1:300) and a rabbit antibody anti-H3K9me3 - Abcam (ab8898, 1:500), overnight at 4 °C. Cells incubated without primary antibodies were used as negative controls for all assays. In the next day, after washing 3x for 10 min each, cells were incubated with secondary antibodies Alexa Fluor 568-conjugated goat anti-rabbit IgG (Life Tech, cat. # A-11036 and Alexa Fluor 488-conjugate goat anti-mouse IgG (Life Tech, cat. #: A-11029) both at RT for 1h. Then, after washing for 3x for 10 min each, samples were mounted on microscope slides and with Prolong Gold Antifade Mountant (Life Tech, cat. # P36935). Images were captured using a confocal microscope (TCS-SP5 AOBS; Leica, Soims, Germany) using laser excitation and emission filters specific for Alexa 488 and Alexa 568. Digital images were analyzed by evaluating each nucleus fluorescent intensity using ImageJ-Fiji image processing software (National Institutes of Health, Bethesda, MD). The fluorescent intensity (average mean gray value) of each channel were measured by manually outlining each nucleus and adjusted against background.

### Protein analysis

All steps for histone acid extraction and western blot were described in details previously, with few modifications ^49^.

Briefly, histones were extracted using the EpiQuick Total Histone Extraction Kit (Epigentek, Farmingdale, NY, USA, cat. # OP-0006) following the manufacturer’s instructions with few modifications. After harvested, cells were centrifuged at 300 xg for 5 min at 4°C, and the pellet were incubated in 200μL of pre-lysis buffer, and centrifuged again at 9,500 xg for 1 min at 4 °C. Then, the supernatant was removed and the pellet was resuspended in 60μL of lysis buffer. After incubation for 30 min at 4 °C, they were centrifuged at 13,500 xg for 5 min at 4 °C. The containing acid-soluble proteins present in the supernatant was transferred into a new vial and neutralized with 0.3 volumes of balance buffer supplemented with DTT. The proteins extracted were quantified by Qubit 2.0 (Life Technologies) and stored in aliquots at −20 °C.

### Western Blotting for quantification of H3K9me2 levels

To measure the relative H3K9me2 levels, 0.5μg of histones were mixed with 4x Laemmli buffer (Bio Rad Laboratories, Hercules, CA, USA cat. #161-0747), following by denaturation at 98 °C for 5 min, and loaded into an acrylamide gel. Proteins were fractionated by size on a 4%–15% SDS-PAGE gel run at a constant 100 V for 90 min. Then, proteins were electroblotted using a semi-dry transfer system (Trans Blot turbo Bio Rad), onto PVDF membranes (Bio-Rad) at a constant 25 V for 3 min. The membranes were blocked with 5% BSA in Tris-buffered saline (TBS) + 0.1% of TWEEN (TBS-T) for 1 h at room temperature, following by incubation overnight at 4 °C under agitation with the primary antibody anti-H3K9me2 (Abcam ab1220) diluted 1:5,000 in TBS-T + 1% BSA solution. In the next day, the membranes were washed 3x in TBS-T for 5 min each and incubated for 1 h with peroxidase-conjugated anti-rabbit secondary antibody (Sigma, cat # A0545) diluted 1:10,000 in TBS-T + 1% BSA solution. The membranes were incubated with Clarity Western ECL Substrate (Bio Rad cat. # 170-5060) for 30 seconds, and images were captured using the ChemiDoc MP Imaging System (Bio-Rad). The analysis of images and band quantification were made using the software Image Lab 5.1 (Bio-Rad). The loading control histone H3 was used to calculate the abundance of H3K9me2. To this end, the same amount of proteins was run in parallel, then the membranes were incubated overnight at 4 °C under agitation with anti-histone H3 antibody (Sigma, cat. # H0164) diluted 1:10,000 in TBS-T + 1% BSA solution, then washed and developed as described above.

### Somatic cell nuclear transfer

SCNT was performed as described ^52^, with few modifications. Bovine ovaries were obtained from a local slaughterhouse and transported to the laboratory in saline solution at 38.5°C. Using a syringe and 18G needle from 3 to 6 mm antral follicles, cumulus oocyte complexes (COCs) were aspirated and cultured in Tissue Culture Medium 199 (TCM199) supplemented with 10% FBS, 0.2 mM pyruvate, 50 μg/mL gentamicin, 0.5 mg/mL FSH, and hCG, 5 U/mL (Vetecor, Hertape Calier) for 18h under mineral oil at 38.5°C and in atmosphere with 5% CO_2_. Then, to remove cumulus cells from oocytes, manual pipetting was made in the presence of hyaluronidase (0.2%). Only oocytes with evident first polar body (PB) were selected for enucleation. To visualize the DNA oocytes were labeled with 10 μg/mL of the DNA fluorochrome Hoechst 33342 for 15 minutes at RT in SOF supplemented with 5□mg/mL BSA, 2.5% FCS, 0.2□mM sodium pyruvate, gentamicin ^53^. Then, they were washed and transferred to a micromanipulation drop of Medium 199, Hanks’ Balanced Salts plus Hepes (Gibco, Grand Island, US, CAT 12350-039) supplemented 10% FBS, 0.2 mM pyruvate, 50 μg/mL gentamicin, and 7.5 μg/ mL cytochalasin B, under mineral oil. All procedures were performed on an inverted microscope (Nikon Eclipse Ti) equipped with Hoffman optics and Narishige micromanipulators (Tokyo, Japan). A 15-um inner diameter enucleation pipette (ES TransferTip; Eppendorf) was used to aspirate and remove the PB and metaphase II plates. The aspirated cytoplasm was exposed to UV light to confirm for the presence of PB and metaphase plate. After enucleation, one single donor cell from control or treatment, at the same cell passage, was placed into the perivitelline space of each enucleated oocyte. To fuse them the reconstructed embryos were placed into a fusion chamber filled with 0.3 M mannitol in a Multiporator (Eppendorf, Hamburg, Germany) with settings of one pulse of alternating current (5 seconds, 50V/cm) followed by two continuous current electric pulses (45 μs each, 1.75 kV/cm). Then, they were incubated for at least 60 min to select the successfully fused. After 26 h post-maturation, the fused SCNT couplets were activated with 5 μM ionomycin in TCM199-HEPES (supplemented with 1 mg/mL fatty acid-free BSA) for 5 minutes and moved into TCM199-HEPES medium (supplemented with 30 mg/mL BSA) for 3 minutes. Then, cultured for 3h in SOF medium supplemented with 6-dimethylaminopurine. Following activation, presumptive zygotes from cell treatment were washed and placed into droplet of 100 μL SOF medium, while for the embryo treatment groups, zygotes from non-treated cells were split into drops of control (SOF medium supplemented with 0.01% of DMSO) or treatment (SOF medium supplemented with 250nM of UNC0638 until reach 8/16 cells stage 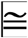 55h) placed in different plates under mineral oil at 38.5°C and in atmosphere with 5% CO_2_. Developmental rates were evaluated at day 2 for cleavage and day 7 for blastocyst. Control for oocyte quality, activation procedure, and in vitro culture was made using activated metaphase II-arrested oocytes, using the same activation protocol described above (parthenogenetic embryos). Embryos at one cell stage (18 hours post-activation), 8/16-cells stage (55 hours post-activation), and at blastocyst stage were collected either for immunostaining procedure or for gene expression analysis.

### Embryo immunostaining

Immunostaining was performed as described previously ^50^, with modifications. Embryos at 1/2-cell stage (18h post activation), 8/16-cells stage (55h post activation), and blastocyst at day 7 after activation were collected and washed in PBS+PVA. Then, embryos were fixed with 4% paraformaldehyde for 15 min and permeabilized with D-PBS + 1% Triton X-100 for 30min. Exclusively for 5mC and 5hmC staining, following permeabilization step, we treated embryos with 4N HCl for 15 min. Next, we proceed with neutralization for 20 min with 100 mM Tris-HCl buffer (pH 8.5). After blocking altogether for 2 h in D-PBS + 0.1%Triton X-100 + 1% BSA, embryos were placed in primary antibody solution consisting of blocking buffer, a mouse antibody anti-H3K9me2 - Abcam (ab1220, 1:300), a rabbit antibody anti-H3K9me3 - Abcam (ab8898, 1:500), a mouse antibody anti-5-mC - Abcam (ab10805, 1:500), and a rabbit antibody anti-5-hydroxymethylcytosine (5-hmC) antibody - Abcam (ab214728, 1:500) overnight at 4 °C. After washing 3x for 10 min and 3x for 20 min each, embryos were incubated with secondary antibodies Alexa Fluor 568-conjugated goat anti-rabbit IgG (Life Tech, cat. # A-11036 and Alexa Fluor 488-conjugate goat anti-mouse IgG (Life Tech, cat. #: A-11029) both at RT for 1h. Then, after washing for 3x for 10 min and 3x for 20 min each, embryos were mounted on slides with Prolong Gold Antifade Mountant (Life Tech, cat. # P36935). Z-stack images were captured using a confocal microscope (TCS-SP5 AOBS; Leica, Soims, Germany) using laser excitation and emission filters specific for Alexa 488 and Alexa 568. Pictures were false-colored using LAS Lite Leica software for informative purpose according to each epigenetic mark (H3K9me2-green, H3K9me3-red, 5mC-cyan, and 5hmC-magenta); no further adjustments or image processing was done. Digital images were analyzed by evaluating each nucleus fluorescent intensity using ImageJ-Fiji image processing software (National Institutes of Health, Bethesda, MD). After maximum projection reconstruction of Z-stacks, the fluorescent intensity (average mean gray value) of each channel were measured by manually outlining each nucleus and adjusted against cytoplasmic background. At least 10 embryos at each stage from 3 different replicates were analyzed.

### RNA extraction, reverse transcription and quantification

Total RNA of 8/16-cell embryos (n= 5) and blastocysts (n =5) from 3 biological replicates were extracted using TRIzol Reagent (InvitrogenTM, Carlsbad, CA) and Linear Acrylamide (InvitrogenTM, Carlsbad, CA), according to the manufacturer’s instruction. Quality and quantification of RNA were performed using NanoDrop 2000 spectrophotometer (Thermo Fisher Scientific; Absorbance 260/280nm ratio). Next, RNA was treated with Deoxyribonuclease I, Amplification Grade (DNaseI, Invitrogen; Carlsbad, CA) for 15min at room temperature to digest any contaminating DNA. DNase I was inactivated by adding 1□L of EDTA in the reaction for 10 min at 65°C. Double-stranded cDNA from 8/16-cell embryos and blastocysts were synthesized from 160ng of total RNA with final volume of 20μl using High Capacity cDNA Reverse transcription Kit (Thermo Fisher). The reaction was composed by total RNA, 10x RT Buffer, 10x RT Random primers, 25x dNTP MIX, nuclease-free water and Reverse Transcriptase. After, the reactions were incubated at 25°C for 10 minutes, 37°C for 120 minutes and 85°C for 5 minutes. Quantitative real-time polymerase chain reactions (qRT-PCR) were performed in QuantStudio 6 Flex (Applied Biosystems) equipment using a final volume of 10 μL per well with 2x Power SYBR Green RT-PCR Master Mix (Applied Biosystems), 1 μM/mL of forward and reverse bovine-specific primers, 6.95ng of cDNA and nuclease-free water. The primers were designed using the software Primer-BLAST (NCBI) based upon sequences available in GenBank (Supplemental Table 1). The primers were also sequenced to test their specificity. *DNTM1, DNMT3A, DNMT3B* and the reference genes PPIA and RPL15 were used as described previously ^54,55^. Amplification were performed with initial denaturation at 95°C for 2 minutes, following by 45 cycles at 95°C for 15 seconds and 60°C for 1 minute, followed by melting curve analysis to verify the presence of a single amplified product. The mRNAs were considered present when the cycle threshold (CT) was less than 37 cycles and presented adequate melting curve. Samples of mature oocytes, 8-16 cells embryos and blastocysts were analyzed in duplicate and the CT data was normalized using geometric mean of PPIA and RPL15 reference genes.

### Statistical analysis

Statistical analysis was performed using GraphPad Prism 7 (GraphPad Software, San Diego, California, USA). Data were tested for normality of residuals and homogeneity of variances using the Shapiro-Wilk test and analyzed as follows. Immunostaining and western blot data were analyzed by Student’s *t*-test. Gene expression experiments data were analyzed by one-way ANOVA followed by Tukey’s post hoc test. Frequency data (developmental rates) were analyzed by Student’s *t*-test. Differences with probabilities of P < 0.05 were considered significant. In the text, values are presented as the means ± the standard error of the mean (S.E.M.). All experiments were repeated at least three times unless otherwise stated.

## Data Availability

The datasets generated during and/or analyzed during the current study are available from the corresponding author on reasonable request.

### Author Contributions

R.V.S. and F.V.M. conceptualized the study; R.V.S. collected the samples; R.V.S., J.R.S., T.H.C.B, D.R.A., M.C., A.B., A.C.F.C.F.A, and C.H.M. performed the experiments; R.V.S., J.C.S., F.P., and M.R.C performed formal data analysis and interpretation; F.V.M., F.F.B., L.J.O, L.C.S, and P.J.R manage and coordinate research planning and execution, and oversight the research activity; R.V.S., L.C.S., M.R.C., and F.V.M. wrote the manuscript with input of all authors. All the authors discussed the results, critically reviewed and edited the manuscript.

## Acknowledgment

The authors would like to thank the staff and students at the LMMD, Jessica Brunhara Cruz, João Vitor Puttini Paixão, and Rodrigo Barreto for their assistance with the sample collections, laboratory procedures and discussions. The slaughterhouses Olhos D’água and Vale do Prata for gently provided ovaries for this study. Rafael Vilar Sampaio is supported by São Paulo Research Foundation - FAPESP, grant number #2015/08807-6 and previously by the grants #2015/25111-5 and #2013/07160-3. This work was granted by São Paulo Research Foundation – FAPESP, grant number #2012/50533-2, #2013/08135-2, #2014/21034-3, #2014/22887-0 and #2014/50947-7; National Counsel of Technological and Scientific Development (CNPq) grant number 465539/2014-9 and CAPES. The funders had no role in study design, data collection and analysis, decision to publish, or preparation of the manuscript.

## Figure Legends

### Supplemental Data Legends

**Supplemental Figure 1** Full immunoblots from data presented on Figure 1. (**A**)-Total histone 3. 1 (**B**) Levels of H3K9me2

**Supplemental table 1**. Primer sequences used in Real Time qPCR.

Accession number Primer Sequences (5’-3’) Product (bp) AT (°C)

