## Supplementary material for "Catalytic inhibition of H3K9me2 writers disturbs epigenetic marks during bovine nuclear reprogramming": Supp Figure 1

A	B

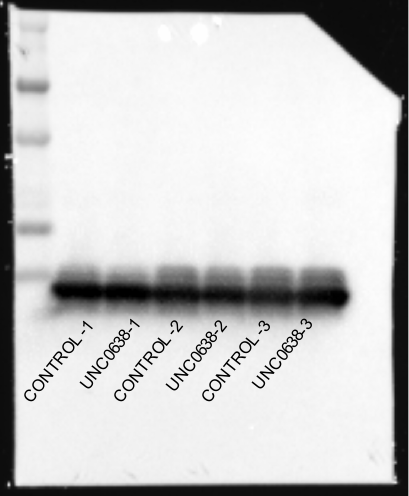

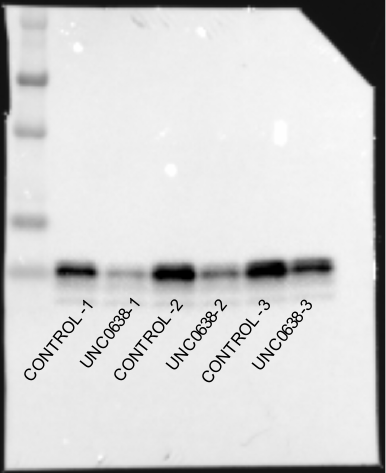

**Supplemental Figure 1** Full immunoblots from data presented on Figure 1. (**A**)-Total histone 3. 1 (**B**) Levels of H3K9me2
